# Spinal Circuit Mechanisms Constrain Therapeutic Windows for ALS Intervention

**DOI:** 10.1101/2025.06.30.662290

**Authors:** Beck Strohmer, Kaitlyn Grosh, Roser Montañana-Rosell, Santiago Mora, Jessica Ausborn, Ilary Allodi

## Abstract

Amyotrophic Lateral Sclerosis (ALS) is a fatal neurodegenerative disease characterized by progressive breakdown of neural circuits which leads to motoneuron death. Earlier work from our lab showed that dysregulation of inhibitory V1 interneurons precedes the degeneration of excitatory V2a interneurons and motoneurons and that stabilizing V1–motoneuron connections improved motor function and saved motoneurons in the SOD1^G93A^ ALS mouse model. However, the optimal timing for this intervention remains unclear. To address this, we developed a spiking neural network model of spinal locomotor circuits to simulate healthy and ALS-like conditions. By modeling changes in network connectivity and synaptic dynamics, we predict that V1 dysregulation induces hyperexcitation in motoneurons which is preferentially observed in flexor motoneurons leading to the disruption of flexor-extensor coordination, and potentially contributing to selective vulnerability of flexor motoneurons. Stabilizing V1 synapses preserved motor output even after motoneuron loss, suggesting that therapeutic benefit is possible into symptomatic stages. However, model predictions also highlighted that after sustained synaptic loss and the development of slower synaptic dynamics within the network, synaptic stabilization leads to maladaptive extensor-biased activity, suggesting that excitatory/inhibitory balance impacts treatment effectiveness. Finally, the model indicated that V1 stabilization could lead to rescue of the V2a excitatory interneurons, a finding that we were able to confirm experimentally in the SOD1^G93A^ ALS mouse model. By exploring different scenarios of synaptic loss and cell dysregulation during synaptic stabilization, our models provide a framework for predicting candidate time windows for spinal circuit interventions, which may guide future preclinical investigations.

## 1. Introduction

Amyotrophic Lateral Sclerosis (ALS) is a debilitating disease characterized by progressive motoneuron death with fatal consequences (13; 33). The exact mechanisms causing motoneuron degeneration in ALS remain elusive, and there is no known cure.

ALS is defined by a loss of motoneurons in the spinal cord and brainstem, in addition to the loss of corticospinal neurons. Functionally, this translates to muscle wasting and progressive paralysis of patients. The progression of ALS is thought to be the consequence of both cell autonomous and cell non-autonomous mechanisms (42). Cell autonomous mechanisms produce aberrant firing of motoneurons through intrinsic neuronal properties while loss of synaptic connectivity within the pre-motor network contributes to cell non-autonomous changes (33).

Approximately 80% of ALS cases are characterized by spinal onset, while the rest show more aggressive bulbar onset (37). In both forms, loss of somatic motoneurons leads to muscle weakness and spasticity, reflecting degeneration of spinal circuits and bulbar (medullary) motor nuclei within the brainstem that control rhythmic motor functions such as walking, breathing, and swallowing (17; 16; 23; 18). The timing of impairment in these functions depends on the onset type (29). In the spinal onset SOD1^G93A^ mouse model, lumbar locomotor central pattern generators (CPGs), the neural circuits that generate the basic locomotor pattern in the lumbar spinal cord, are affected early in disease (2; 32; 41).

The degeneration of the lumbar locomotor CPGs is characterized by synaptic loss (2; 32; 34; 33; 41; 35) and dysregulation of key ventral interneu-rons in the SOD1^G93A^ ALS mouse model (2; 32; 34; 33). This dysregulation contributes to motor impairments, since ventral interneuron classes play a central role in locomotor pattern generation. These classes can be defined by their transcription factor expression, neurotransmitter phenotype, and connectivity patterns, and each contributes distinct functions to the locomotor network. Inhibitory V1, V2b, V0d, and excitatory V0v, V0c, V0g, and V2a populations comprise, among others, the core of the CPG (23; 18). V1 populations include Ia interneurons that provide reciprocal inhibition to antagonistic motor pools, Renshaw cells (RCs) that provide recurrent inhibition to motoneurons, and other V1 interneurons with diverse functions, including a population we refer to as V1*_RG_* that affects locomotor frequency and contributes to flexor-extensor alternation together with V2b interneurons (15; 47; 10). The excitatory populations V0c and V2a, both increase activation of ipsilateral motoneurons (10).

Despite experimental advances in identifying CPG neuronal subtypes, their synaptic connectivity remains incompletely understood (39; 18). While *in vivo* and *ex vivo* studies have shed light on the roles of individual populations in locomotion, probing their activity while simultaneously assessing ALS-related degenerative dynamics, remains challenging (32). To address these issues, we developed the first computational model of ALS disease progression in spinal microcircuits, linking cell-type specific dysregulation to motor output.

We implemented the spinal locomotor CPG for flexor-extensor alternation as a spiking neural network, incorporating relevant interneuron and motoneuron populations (MNPs). The model implements a flexor-driven rhythm generator architecture, an organization supported by experimental evidence that flexor circuits are the primary drivers of locomotor rhythm (11; 40; 6). In this framework, both the flexor and extensor are able to generate bursting activity that alternates due to reciprocal inhibition (18). However, flexor bursting is more readily induced (19). Moreover, during fictive locomotion in isolated spinal cord preparations, spontaneous deletions of flexor bursts result in tonic extensor activity, whereas deletions of extensor bursts do not disrupt the ongoing flexor rhythm (48). These observations support a flexor-driven, rather than symmetric, organization (43; 40).

We used motor phenotypes of healthy and SOD1^G93A^ ALS mouse models to validate network connectivity and subsequent dysregulation (2). Degeneration in the model was dictated by the documented pattern of degeneration observed in the CPG network in SOD1^G93A^ mice. Motoneurons and V1, V0c, and V2a interneurons undergo dysregulation and/or connectivity changes at different stages of ALS progression (41; 2; 32). V1 synaptic projections are lost early, followed by the loss of V1-specific markers (e.g., Engrailed 1 transcription factor) and cell death, preceding motoneuron and V2a interneuron loss (2; 32). In contrast, V0c interneurons respond to network degeneration by temporarily increasing the size of their projecting synapses (38; 20; 41; 7). Beyond these network changes, recent work by (35) found that inhibitory and excitatory postsynaptic currents between RCs and motoneurons showed increased rise and decay times early in disease. These altered synaptic dynamics are later offset by homeostatic compensation, though still at a pre-symptomatic stage, returning rise and decay times to healthy values.

At the behavioral level, V1 dysregulation results in a decrease in locomotor speed and stride length, and hindlimb hyperflexion during early stages of disease in SOD1^G93A^ mice (2; 34) (Fig. 1A). Hindlimb hyperflexion has been repeatedly linked to V1 interneuron loss in mouse models (47; 9). V1 interneurons have also been reported to provide stronger connectivity to flexor motoneurons (9) and differences in the rate of degeneration of flexor and extensor muscles have been observed in patients (30). Together, this raises the question if motoneurons controlling flexor and extensor muscles are affected at different rates.

**Figure 1:**
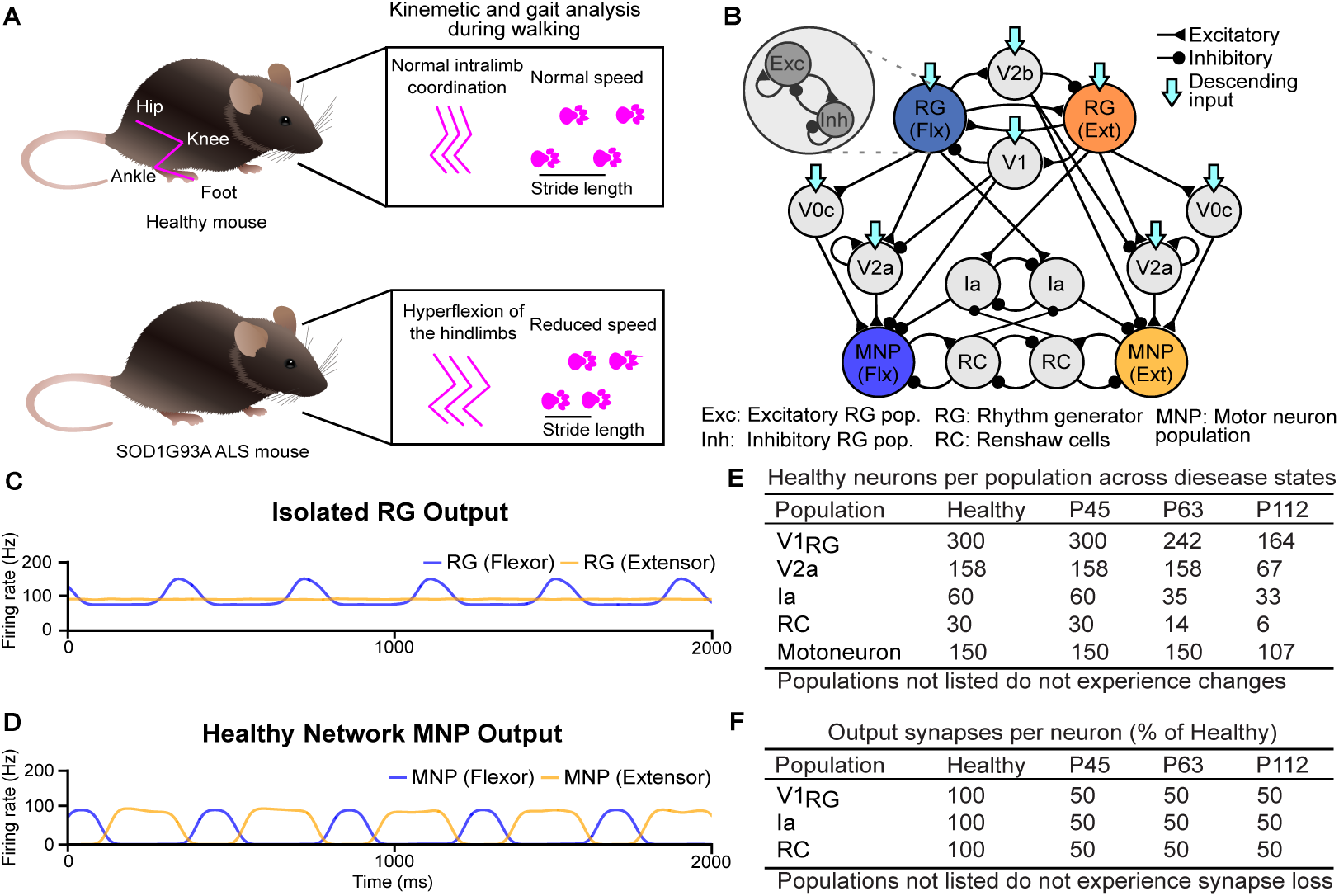
Computational model framework and baseline network properties. **A**) Early motor phenotype changes observed in the SOD1^G93A^ mouse model. **B**) Connectivity diagram of the healthy network model. **C**) Healthy output of the isolated rhythm-generating populations reflecting the flexor-driven architecture (43) with the flexor in bursting and the extensor in tonic mode. **D**) Healthy MNP output of the complete network showing alternating flexor-extensor activity with an extensor-dominated step cycle. **E**) Number of neurons modeled per time point, replicating biological ratios. **F**) Percentage of synaptic input loss per V1 subpopulation and disease state.(32).

Further work conducted in our laboratory demonstrated that V1-restricted overexpression of the extended synaptotagmin 1 (Esyt1) presynaptic organizer led to stabilization of V1 interneuron synapses onto motoneurons, an increase in the number of spared motoneurons, and ameliorated motor functions in the SOD1^G93A^ mouse model (34). Esyt1 is a calcium dependent protein that facilitates endoplasmic reticulum and plasma membrane tethering as well as vesicle trafficking (25), and is preferentially found in neurons resistant to ALS (3). These findings suggest a therapeutic potential of Esyt1, but the optimal time window and E/I balance between excitation (E) and inhibition (I) for intervention, remain elusive. In our study, E/I balance refers to the relative contributions of excitation and inhibition through the number and strength of synapses in the network exciting or inhibiting their postsynaptic targets.

We therefore set out to address these questions using computational modeling, which enabled systematic investigation of the effects of synaptic stabilization interventions by comparing the output of the treated network with that of the degenerated state across different disease stages. We modeled healthy locomotor output using a flexor-driven CPG, and then simulated ALS-related degeneration to capture flexor hyperexcitation. Using this framework, we tested how sparing V2a interneurons or motoneurons altered network dynamics, and whether V1-targeted synaptic stabilization might improve motoneuron output, promote survival of vulnerable interneurons, and restore motor function. Importantly, we then evaluated a key model prediction experimentally, focusing on interneuron survival and synaptic input changes.

Overall, by applying known degeneration and intervention patterns, our models identified a probable time window for effective treatment with synaptic stabilization.

### 1.1. Results

#### 1.1.1. Healthy network output simulated via a flexor-driven CPG architecture

We compared outputs of progressively degenerating spinal CPG networks using computational models to recapitulate motor phenotypes observed in healthy and SOD1^G93A^ mice (Fig. 1B,2A-C). Healthy output was assessed in both the isolated rhythm generator (RG) populations (Fig. 1C) and the flexor and extensor Motoneuron Population (MNP) activity of the full net-work (Fig. 1D).

**Figure 2:**
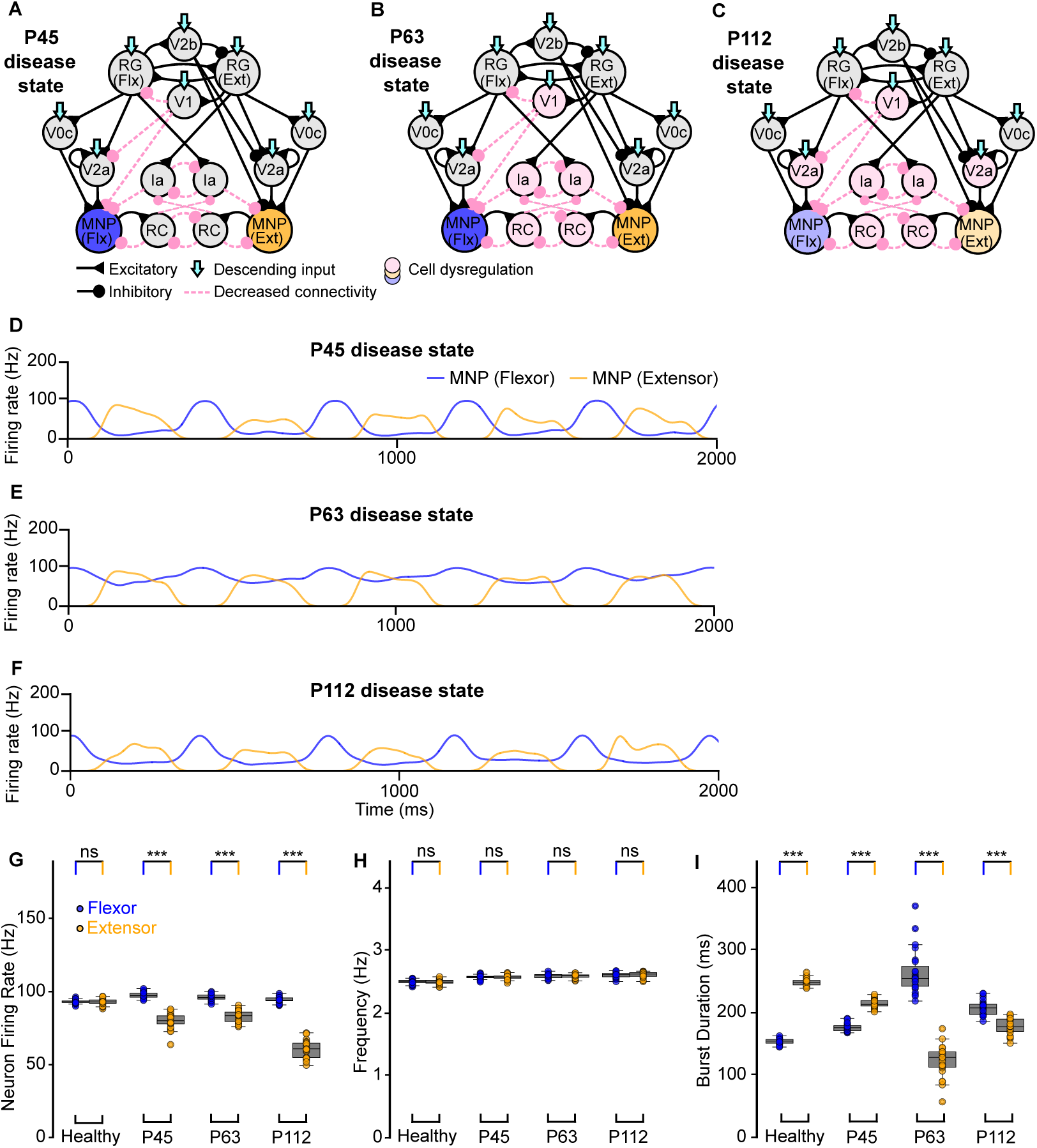
Computational models of the locomotor CPG during ALS degeneration. **A**-**C**) Connectivity diagrams of the network without intervention showing synaptic and cell dysregulation across disease stages. **D**-**F**) MNP output at each time point without intervention. **G**-**I**) Variance plots comparing flexor versus extensor activity showing flexor-biased activity across disease time points **(G)**, similar flexor-extensor frequency within time points **(H)**, and an extensor-dominated step cycle in early disease **(I)**. Variance plots indicate the 50% median line with box edges representing the first and third quartiles, and whiskers at 1.5 times the interquartile range. Significance comparing flexor versus extensor was assessed using a Wilcoxon Signed-Rank test, N=25; p *<*= 0.001 = ***, 0.001 > p *<*= 0.01 = **, 0.01 > p *<*= 0.05: *, p > 0.05 = ns.

A tonic excitatory drive to the RG populations was introduced to model brainstem drive and tuned to reproduce physiologically realistic oscillation frequencies of 2-3Hz (Fig. 1D, 2H, Fig. B.11A) and sparse firing rates in the flexor and extensor MNPs (Fig. 2G, Fig. B.11A). The healthy network model also generated alternating flexor-extensor output (Fig. B.11A) with an extensor-dominated step cycle (Fig. 2I, Fig. B.11A). An extensor- (flexor-) dominated step cycle is defined by longer extensor (flexor) MNP bursts and should be distinguished from extensor- (flexor-) biased activity, where one MNP fires at a higher rate than the other. This behavior is consistent with previous modeling and experimental observations (43; 9; 47). In the flexor-driven architecture, the intrinsically bursting flexor sets the rhythm, while the extensor is tonically active between flexor bursts. This organization naturally produces an extensor-dominated step cycle at low to intermediate locomotor frequencies, recapitulating experimental network dynamics (43; 40).

Together, this model establishes a validated healthy baseline that serves as the reference point for subsequent simulations of ALS-related degeneration.

#### 1.1.2. Flexor motoneurons exhibit hyperexcitation after loss of inhibition

After validation of the healthy network model, disease progression was introduced according to previously described network changes in SOD1^G93A^ mice: early loss of V1 synaptic projections at postnatal day (P) 45 (2), dysregulation of V1 populations at P63 (2; 32), and further exacerbation of V1 dysregulation together with dysregulation of motoneurons and V2a interneurons at P112 (32). These stages were implemented by removing cells and connections in proportions derived from experimental data (Fig. 1E) (32), 2A-C).

The sample traces in Figure 2D-F show changes in flexor-extensor outputs across three disease stages, resulting from the above described degenerative changes. Flexor motoneuron firing rates increased at all disease time points in comparison to the healthy condition, indicating a persistent shift towards hyperexcitation (Fig. 2G-I, Fig. B.11A). Flexor-biased activity was also observed across all disease time points (Fig. 2G, Fig. B.11A). Moreover, output frequency gradually increased (Fig. 2H, Fig. B.11A), while the step cycle transitioned from extensor-dominated at P45 to flexor-dominated at P63 and P112 (Fig. 2I, Fig. B.11A). Finally, flexor-extensor phase difference was most impacted at P63 (Fig. B.11A). Together, these results confirm that the models reproduce key functional signatures of spinal circuit dysregulation in ALS and provide a framework to explore different intervention strategies.

#### 1.1.3. Stabilizing V2a populations increases output variability after V1 dysregulation

To disentangle the relative contributions of interneuron versus motoneuron survival to network stability, we next focused on the P112 time point and selectively spared either V2a interneurons or motoneurons, or both (3A-C).

The sample traces in Figure 3D, F, H illustrate that flexor-biased activity persisted despite sparing either the V2a populations, motoneurons, or both. When motoneurons were spared but V2a interneurons were lost, the disparity between flexor and extensor firing rates increased (Fig. 3E, Fig. B.11B), indicating that V2a-derived excitation is essential for balancing motoneuron activity. By contrast, sparing V2a interneurons after V1 dysregulation increased variability in burst duration and phase (Fig. 3G, I, Fig. B.11B), compared to stabilizing motoneurons alone (Fig. 3E). This suggests that V2a interneuron survival can stabilize flexor-extensor firing rate but destabilize rhythmic timing. Together, these results predict that V2a interneurons exert a stronger influence than motoneurons on shaping both the E/I balance within the nextwork and the timing of locomotor output, further supporting their pivotal role in ALS-related network dysfunction.

**Figure 3:**
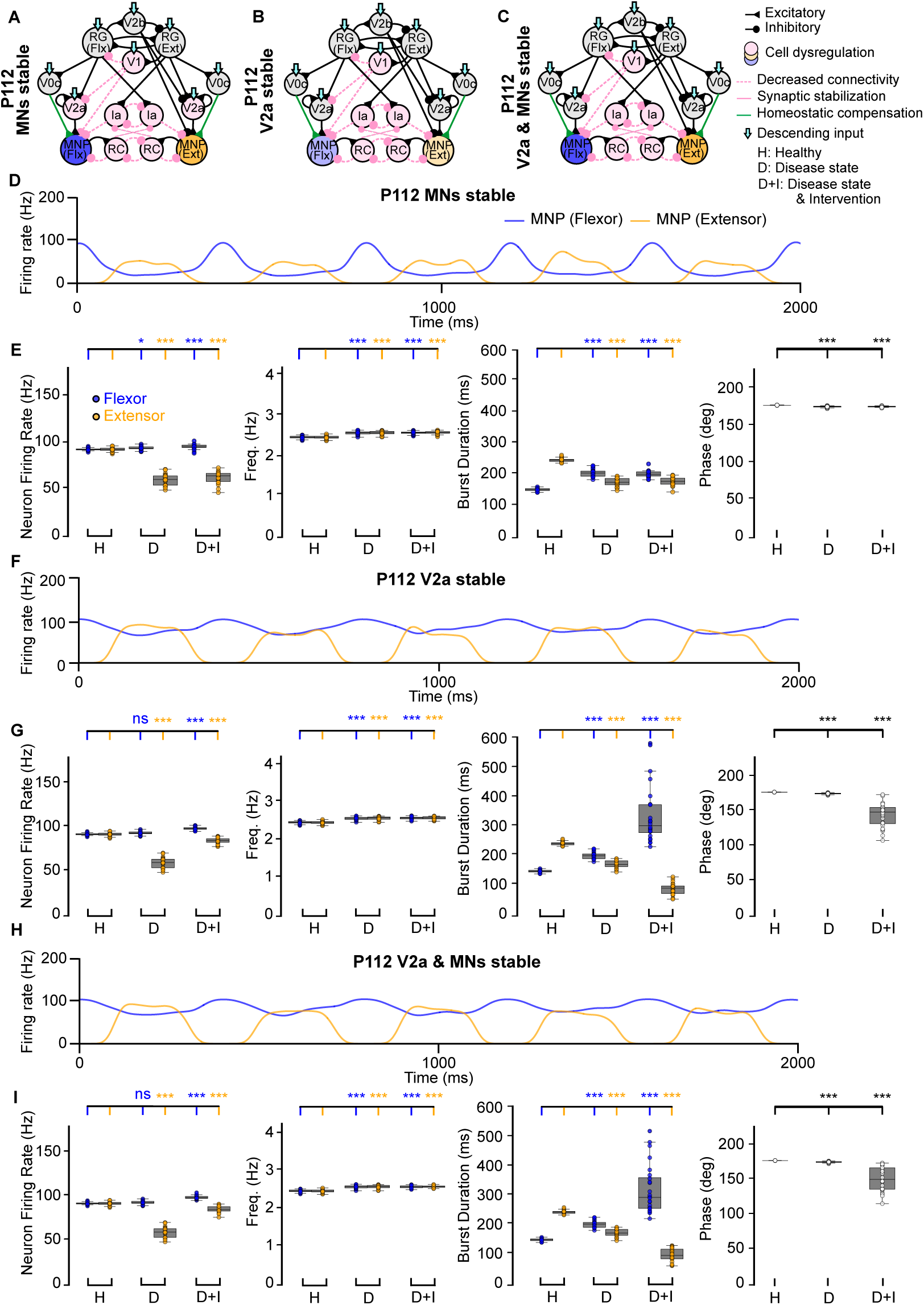
Computational models of the locomotor CPG when stabilizing V2a interneurons and motoneurons after network degeneration. **A-C**) Connectivity diagrams showing network changes and interventions. **D, F, H**) Example outputs of flexor and extensor motoneuron populations. **E, G, I**) Variance plots comparing healthy, disease, and intervention conditions. The loss of V2a cells without intervention increases variability of flexor burst duration as well as phase **(G, I)**. Variance plots indicate the 50% median line with box edges representing the first and third quartiles, and whiskers at 1.5 times the interquartile range. Significance across conditions was assessed using a Kruskal-Wallis H-test with Dunn’s test post hoc, N=25; p *<*= 0.001 = ***, 0.001 > p *<*= 0.01 = **, 0.01 > p *<*= 0.05: *, p > 0.05 = ns.

#### 1.1.4. Exogenous synaptic intervention restores motor output characteristics late in disease

We next simulated synaptic stabilization as a therapeutic strategy to counteract network dysregulation. In our prior work using the ALS mouse model, synaptic stabilization was administered early during disease progression (P45), preceding motoneuron degeneration and exacerbation of network alterations (34). Here, we examined the optimal timing of intervention by strengthening remaining V1 synaptic connections in the model (Fig. 4A-C).

**Figure 4:**
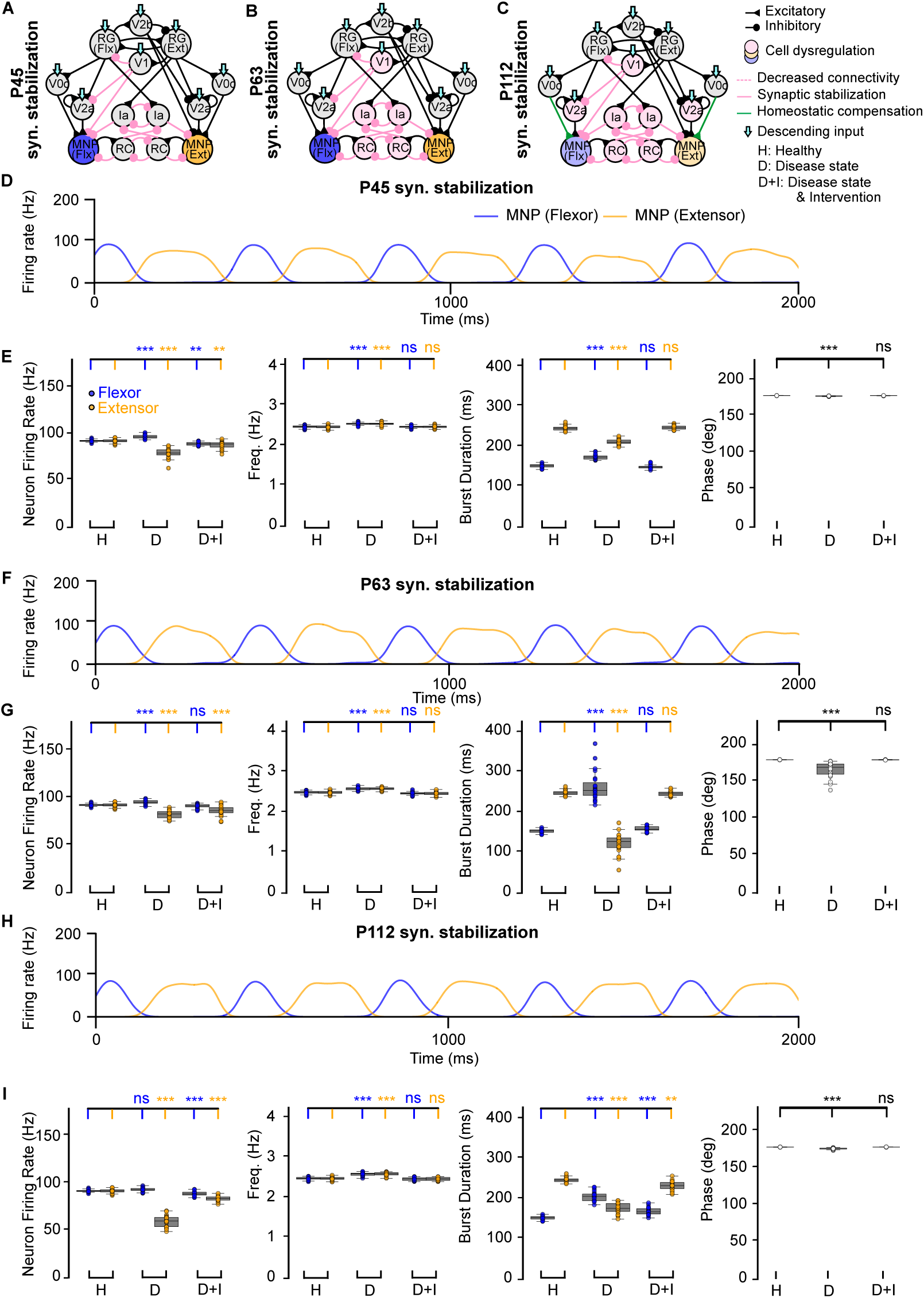
Computational models of the locomotor CPG after synaptic stabilization during ALS degeneration. **A-C**) Connectivity diagrams showing network changes and interventions. **D, F, H**) Example outputs of flexor and extensor motoneuron populations. **E, G, I**) Variance plots comparing healthy, disease, and intervention conditions at different disease states. Synaptic stabilization reduces flexor-biased activity throughout degeneration and restores frequency and phase. Variance plots indicate the 50% median line with box edges representing the first and third quartiles, and whiskers at 1.5 times the interquartile range. Significance across conditions was assessed using a Kruskal-Wallis H-test with Dunn’s test post hoc, N=25; p *<*= 0.001 = ***, 0.001 > p *<*= 0.01 = **, 0.01 > p *<*= 0.05: *, p > 0.05 = ns.

After exogenous intervention via synaptic stabilization, we found a reduction in flexor firing rate and flexor-biased activity at all disease time points (Fig. 4D-I, Fig. B.11C). In addition, frequency and phase were rescued across all time points (Fig. 4E, G, I). Together, these results suggest that synaptic stabilization of V1 projections may ameliorate hyperexcitation of flexor motoneurons during the initial stages of V1 dysregulation, indicating a potential mechanism for further preclinical testing.

#### 1.1.5. Computational models predict firing rate restoration after V1 synaptic stabilization by sparing motoneurons and V2a interneurons

To assess the circuit-level effects of V1 stabilization, we next examined its impact at P112, a symptomatic stage, by selectively sparing V2a interneurons, motoneurons, or both (Fig. 5A-C). Synaptic stabilization performed in the presence of spared V2a interneurons and/or motoneurons showed a return to an extensor-dominated step cycle and restoration of frequency and phase (Fig. 5E, G, I, Fig. B.11D). The sample traces in Figure 5D, F, H qualitatively show that the output improved when compared to the untreated network at P112 (Fig. 2F). Most notably, firing rate also improved when the V2a populations were stable (Fig. 5G, I, Fig. B.11D) and aberrant flexor-biased activity was eliminated when both the V2a interneurons and motoneurons were spared (Fig. 5I, Fig. B.11D). Our model results are consistent with the earlier in vivo study (34) showing that hyperflexion is alleviated after V1 stabilization. In that study, they find that motoneurons are spared, and our model suggests that V2a interneurons may also be spared by such an intervention.

**Figure 5:**
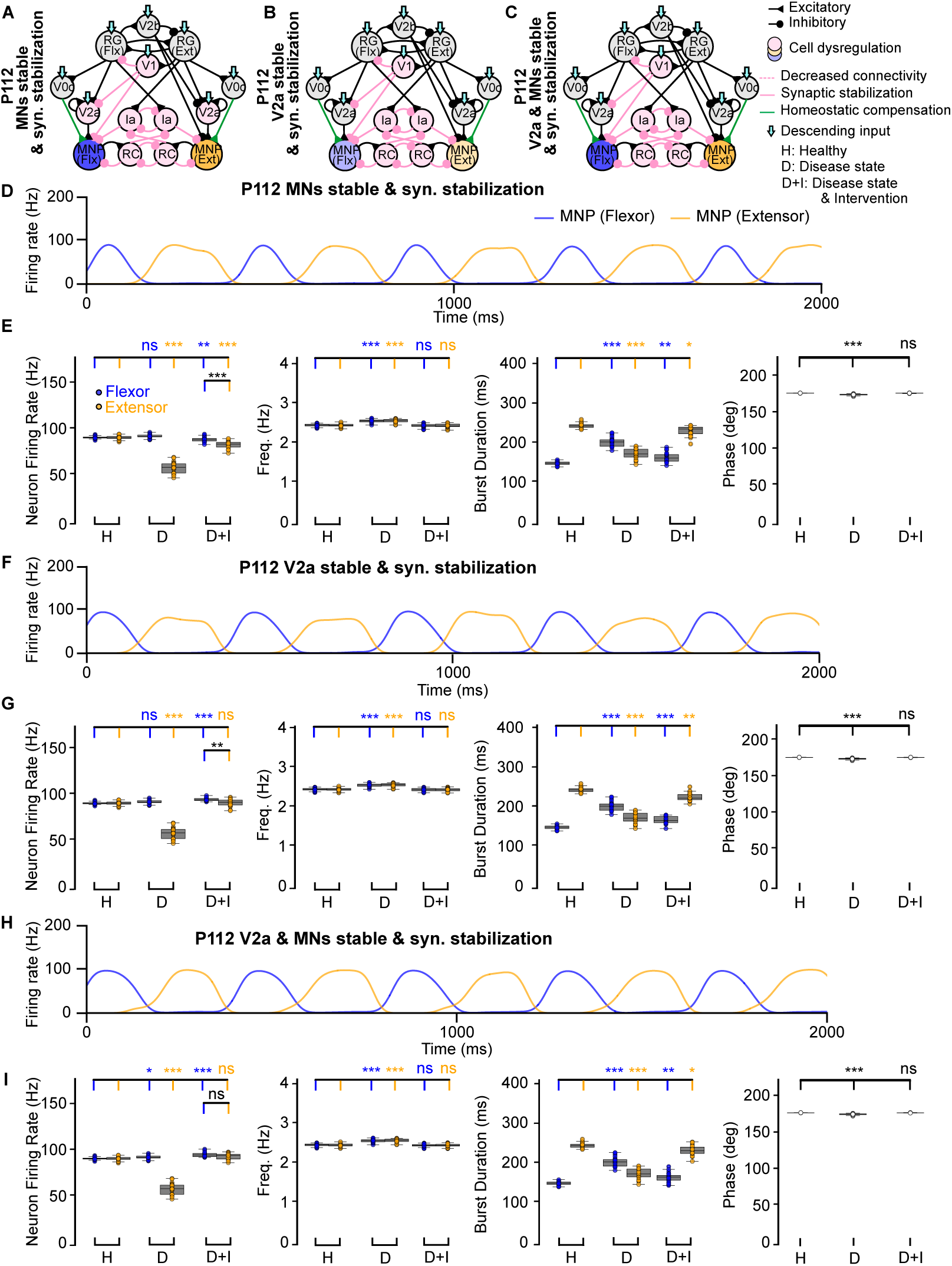
Computational models of the locomotor CPG stabilizing V2a interneurons and motoneurons after synaptic stabilization intervention. **A-C**) Connectivity diagrams showing network changes and interventions. **D, F, H**) Example outputs of flexor and extensor motoneuron populations. **E, G, I**) Variance plots comparing healthy, disease, and intervention conditions. Synaptic stabilization reduces flexor-biased activity and restores the extensor-dominated step cycle, frequency, and phase. Variance plots indicate the 50% median line with box edges representing the first and third quartiles, and whiskers at 1.5 times the interquartile range. Significance comparing flexor versus extensor was assessed using a Wilcoxon Signed-Rank test, significance across conditions was assessed using a Kruskal-Wallis H-test with Dunn’s test post hoc, N=25; p *<*= 0.001 = ***, 0.001 > p *<*= 0.01 = **, 0.01 > p *<*= 0.05: *, p > 0.05 = ns.

To further investigate these results, we also explored V1 synapse and cell rescue in the presence of V2a interneuron and motoneuron loss (Fig. B.8A). In this scenario, with V1 cells and synapses spared, our model also predicted a reduction in flexor-biased activity and restored output frequency and phase difference (Fig. B.8B, C, D). Together, these results suggest that V1 synaptic stabilization alone is insufficient to restore flexor-extensor firing rate and requires sparing of motoneurons and V2a interneurons. This aligns with our previous findings (34), which showed that V1-restricted *ESYT1* overexpression improved survival of motoneurons in the SOD1^G93A^ mouse model. However, the impact of this treatment on other spinal neurons receiving inputs from V1 interneurons (i.e. V2a interneurons) is still unknown.

#### 1.1.6. ESYT1-driven synaptic stabilization increases V2a interneuron survival and VGlut2 positive synapses

While V1 stabilization alone reduced flexor hyperexcitation (Fig. 4), the model predicted that robust restoration of balanced locomotor output required sparing of V2a interneurons (Fig. 5). This led us to hypothesize that V1 stabilization, achieved by overexpression of *ESYT1* administrated through gene therapy, indirectly promotes V2a interneuron survival. Hence, the effects of *ESYT1* overexpression on the V2a interneurons were investigated in the SOD1^G93A^ mice. Firstly, the SOD1^G93A^ mice were crossed with mice expressing cre-lox recombinase under the Engrailed 1 (En1) promoter to allow for V1-restricted *ESYT1* overexpression in the ALS mice. En1 is the specific marker for V1 interneurons (15). Secondly, *ESYT1* overexpression was achieved by intravenous administration of an AAV-PHP.eB cre-dependent vector (34). AAV administration was performed between P28 and P35, double transgenic SOD1^G93A^;En1^cre^ mice received an AAV-PHP.eB-hSYN-DIO-hESYT1-W3SL injection, while control/untreated SOD1^G93A^ mice were injected with an AAV-PHP.eB-hSYN-eGFP-W3SL over-expressing eGFP. The treated and untreated mice were kept until humane end point (e.g., 15% weight loss or paralysis of one limb) and then euthanized for tissue collection and anatomical analysis.

To quantify V2a interneurons, lumbar spinal cord sections between L1-L3 were analyzed using immunohistochemistry for the Chx10 transcription factor, which is a specific marker for V2a interneurons (1; 28) (Fig. 6A-D). Chx10-positive neurons are found in the intermediate area of the lumbar spinal cord (Fig. 6A) and the number of Chx10 neurons in the untreated and treated mice was compared (Fig. 6B-D). Quantifications showed a higher number of Chx10-positive neurons in the *ESYT1* treated mice (t-test, P = 0.0119, t = 3.859, df = 5, untreated N = 3 and treated N = 4), which supports the prediction obtained from our computational model that V2a interneurons are spared together with motoneurons. Moreover, VGlut2 positive synaptic inputs were quantified within the ventral horn of the spinal cord (Fig. 6E). The mean area of pixels positive for VGlut2 was quantified for untreated and treated mice (Fig. 6F-H) and suggested higher mean area values in the *ESYT1* treated mice (t-test, P = 0.0589, t = 2.253, df = 7, untreated N = 5 and treated N = 4) (Fig. 6H).

**Figure 6:**
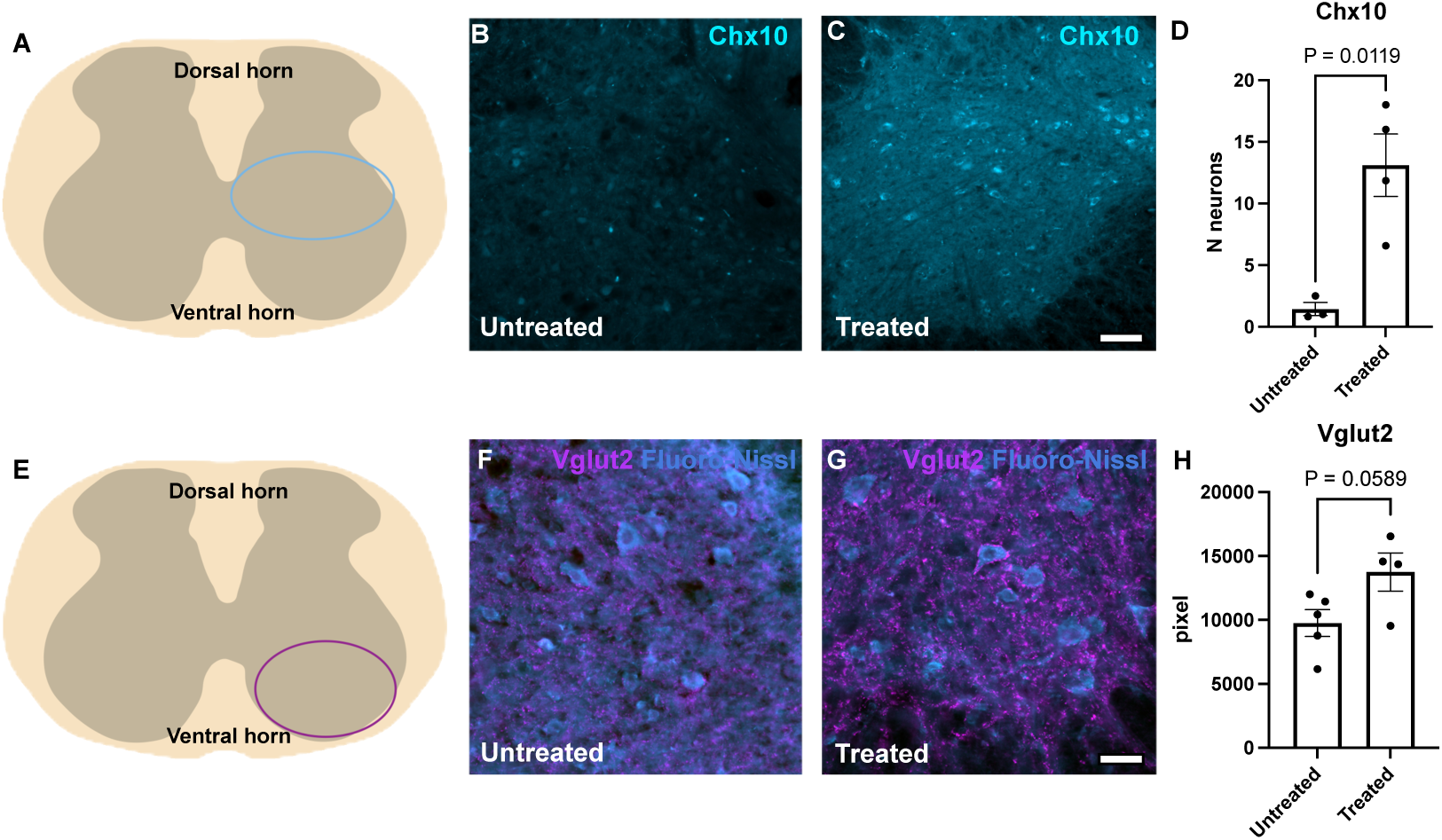
V2a interneuron rescue after synaptic stabilization in the SOD1^ee3A^ mouse model. **A**) Cartoon of a spinal cord cross section. The light blue oval indicates the intermediate region of the spinal cord where Chx10-positive neurons were quantified.**B-C**) Microphotographs show Chx10 expression in the untreated and treated end-stage SOD1^G93A^ mice, respectively. Scale bar in **C**=100 *µ*m, Chx10 staining in cyan. **D**) Variance plot reports increased number of Chx10-positive neurons in the Esyt1 Treated mice compared to the Untreated (unpaired t test, P=0.0119, t=3.859, df=5, untreated N=3, treated N=4). **E**) Purple oval indicates the ventral spinal cord region where VGlut2 positive excitatory synapses were quantified. **F-G**) Microphotographs show VGlut2 staining (magenta) and Fluoro-Nissl (cyan) in untreated and treated end-stage SOD1^G93A^ mice. Scale bar in **G**=50 *µ*m. **H**) Variance plot shows area of pixels quantified in the ventral horn of untreated and treated mice (unpaired t test, P=0.0589, t=2.253, df=7, Untreated N=5, Treated N=4).

#### 1.1.7. Synaptic stabilization is e!ective only after homeostatic restoration of synaptic dynamics

To predict an optimal therapeutic time window, we investigated additional potential restrictions to the timing of synaptic intervention. In SOD1^G93A^ mice, inhibitory synapses from RCs to motoneurons and excitatory synapses from motoneurons to RCs exhibit slower rise and decay times until P48, when endogenous homeostatic compensation restores synaptic dynamics (35). We added these slower synaptic dynamics in our P45 model to the inhibitory connections from RCs and Ias to MNPs and to the excitatory connections from MNPs to RCs to test their effects after loss of V1 synaptic connections in the untreated (Fig. 7A) and treated state (Fig. 7B). These slower synaptic dynamics were only applied to the flexor-side of the model. When we tested applying slower synaptic dynamics to only the extensor-side or symmetrically (Fig. B.9A, B), the flexor-biased activity was exacerbated and the variance of the frequency increased (Fig. B.9C-G). The hyperflexion locomotor phenotype emerges only after P45 in vivo (2). Therefore, we selected the flexor-side slow synaptic dynamics condition, which produced the smallest difference between flexor and extensor firing rates, as the most consistent with the biological condition for further simulations.

**Figure 7:**
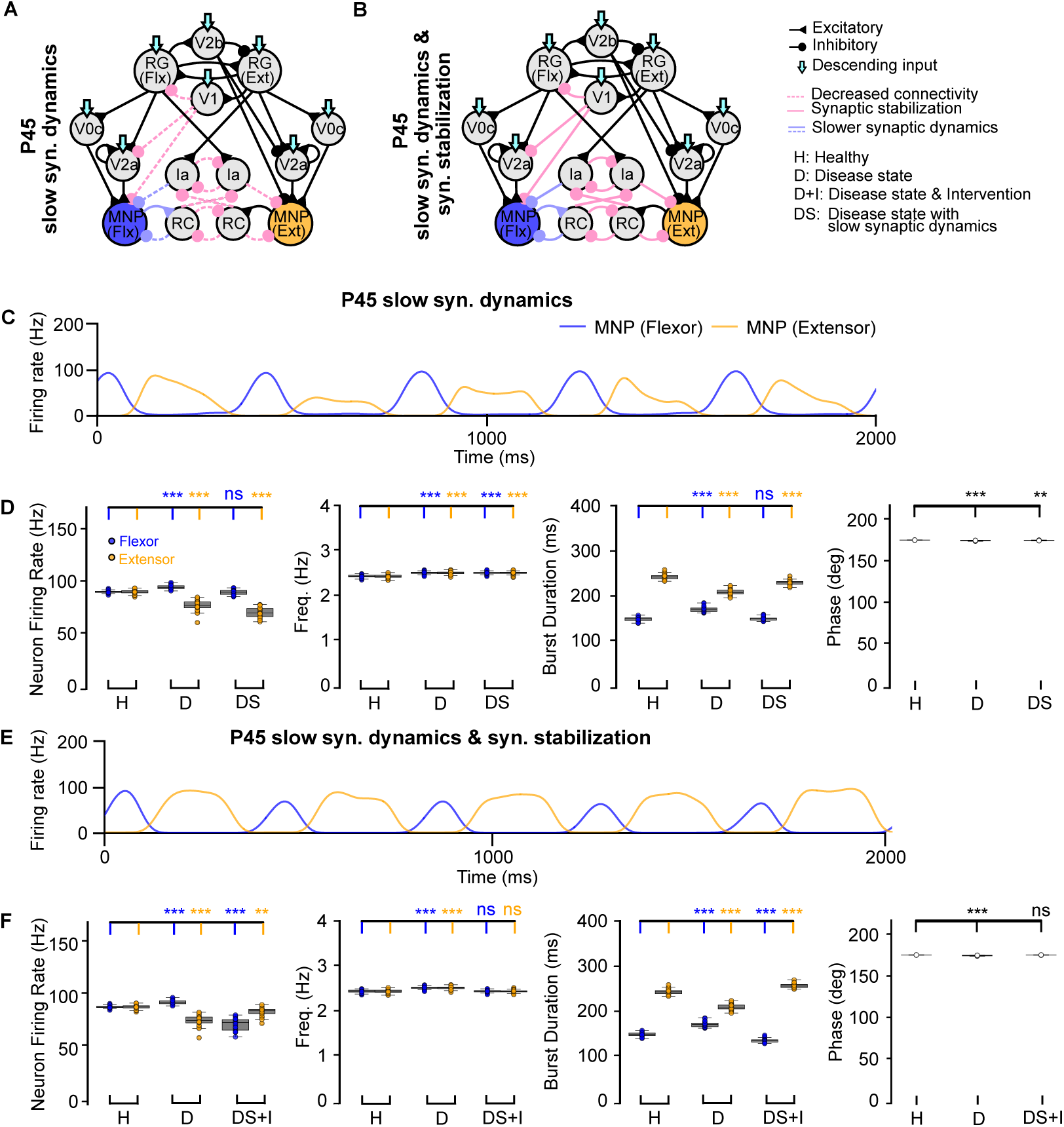
Computational models of the locomotor CPG with and without synaptic stabilization. **A-B**) Connectivity diagrams showing network changes and interventions. **C, E**) Example outputs of flexor and extensor motoneuron populations. **D, F**) Variance plots comparing healthy vs P45 no intervention and slow synaptic dynamics. Stabilization promotes extensor-biased activity during slow synaptic dynamics but restores frequency and phase. Variance plots indicate the 50% median line with box edges representing the first and third quartiles, and whiskers at 1.5 times the interquartile range. Significance across conditions was assessed using a Kruskal-Wallis H-test with Dunn’s test post hoc, N=25; p *<*= 0.001 = ***, 0.001 > p *<*= 0.01 = **, 0.01 > p *<*= 0.05: *, p > 0.05 = ns.

Aberrant flexor-biased activity was observed during slow synaptic dynamics in the untreated state, although flexor motoneuron firing rates remained similar to those in the healthy network (Fig. 7C, D, Fig. B.11E). Interestingly, exogenous synaptic stabilization applied during slow synaptic dynamics induced a shift to extensor-biased activity (Fig. 7E, F, Fig. B.11E). We predict that application of stabilization during this stage would therefore result in maladaptive hyper-extension of the limb, which differs from the hyperflexion locomotor phenotype observed in the SOD1^G93A^ ALS mice. This reinforces the earlier observation that E/I balance plays a role in the effectiveness of stabilization strategies and leads us to suggest that the most optimal window of intervention occurs after homeostatic compensation restores synaptic dynamics.

### 1.2. Discussion

The goal of this study was to determine whether synaptic stabilization of spinal inhibitory circuits could mitigate ALS-related network dysfunction and to predict when such interventions are most effective. By integrating spiking network models with experimental data, we predict that synaptic stabilization of V1 interneurons can reduce flexor-biased activity and preserve motoneuron output, but only after synaptic dynamics have recovered. These findings highlight the importance of intervention timing, reveal mechanisms that may underlie maladaptive responses to premature stabilization, and provide a framework for interpreting flexor-dominant vulnerability observed in ALS patients (30; 31; 21).

This stabilization not only improves motoneuron output, but has also been shown previously to protect motoneurons against dysregulation (34). Building on this and other experimental data (2; 32), our models predicted that V1 stabilization reduces flexor hyperexcitation but that robust restoration of balanced output depends on additional sparing of V2a interneurons. We tested this prediction in the SOD1^G93A^ ALS mouse model and now also demonstrate that V1-restricted stabilization also preserves V2a interneuron populations.

The models also reinforce that there is a limit to how much the original balance in the CPG network can be altered before intervention strategies risk causing maladaptive responses. If synaptic stabilization is applied prior to recovery of synaptic dynamics, the network skews toward extensor-biased activity. This likely arises because prolonging rise and decay times on the flexor side reduces the temporal precision of inhibitory feedback from RCs and Ia interneurons, weakening flexor control and allowing extensor activity to dominate. Thus, although the slower synaptic dynamics were applied asymmetrically to the flexor side, their emergent effect was to shift the balance toward extension. This predicts that early synaptic changes may preferentially affect flexor motoneurons, consistent with recent clinical findings of greater flexor muscle involvement in early-stage ALS (30). In our simulations, this bias arises from the well-established flexor-driven architecture. However, while we grounded our predictions in a flexor-driven architecture based on current available evidence in the literature (11; 40; 6; 19; 48), this is only one of several plausible configurations for locomotor CPGs. Other proposed concepts such as balanced half-center or Unit Burst Generator models (23; 18; 40) may yield different network responses to asymmetric degeneration. Future work could test under what conditions timing-dependent intervention effects are robust across different CPG architectures or instead dependent on specific assumptions about the network architecture. In turn, an experimental confirmation of flexor motoneuron vulnerability in the SOD1^G93A^ mouse model would provide further support of the flexor-driven architecture hypothesis.

From a therapeutic perspective, these findings suggest that interventions targeting synaptic dynamics may need to be temporally tailored. Specifically, it will be important to test whether synaptic dynamics could be a target for treatment. For example, if exogenous synaptic stabilization results in hyperextension, our model indicates that shortening the rise and decay times of synaptic projections onto flexor-side RCs and motoneurons could help restore motoneuron firing rates.

Our model suggests that the untreated network shows larger variation of both burst duration and phase at P63 as compared to P112 (Table B.11). This could be a consequence of how burst duration and phase are calculated, using the intersection of flexor and extensor output signals to determine each. One solution would be to negatively offset the flexor output so that the intersection between signals was more consistent. However, we did not apply this solution in order to keep statistics comparable across disease time points.

Several limitations of the present model must be considered. While our model has recurrent connections within the architecture, feedback from external muscles is not implemented. Therefore, network output cannot fully predict muscle activity. In our results, we show that synaptic stabilization preserves motoneuron firing rates. However, this does not directly translate to healthy muscle activation after motoneuron loss, since fewer motoneurons would be available to innervate muscle fibers. Future studies will incorporate Ia afferent feedback, as this is known to be affected during ALS progression in the SOD1^G93A^ mouse model (4; 33; 35).

Finally, our modeling provides mechanistic insights into frequency changes observed during ALS progression. Our findings show an increase in frequency over progressive disease time points as V1 cells are lost. This is a consequence of our flexor-driven architecture, where the flexor RG serves as the main driver for network oscillations. As the V1*_RG_* population loses projections and cells, it allows the flexor RG to spike more, thereby increasing network frequency. This is similar to results seen in the neonatal mouse hemicord preparation, where hyperpolarization of V1 interneurons increases oscillation frequency (12). By contrast, in the intact spinal cord preparation, hyperpolarization of V1 interneurons decreases oscillation frequency (12). This is consistent with our earlier in vivo study showing a decrease in frequency and stride length at progressive disease time points in the SOD1^G93A^ mouse model (2). Contralateral inhibition has been hypothesized to contribute to this decrease, thereby mediating different outcomes in the hemicord versus full cord; this hypothesis will be addressed in future studies.

## 2. Methods

### 2.1. Neuron Model

This study uses an adaptive exponential integrate-and-fire (AdEx) neuron model to maintain computational efficiency while being able to produce both tonic and bursting firing behaviors. The membrane potential *V_m_*for the neuron at each time step (0.1ms) is calculated according to Equations (1) and (2).

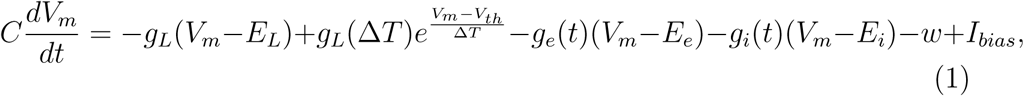

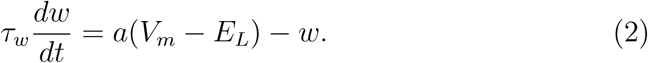

Reset equation:

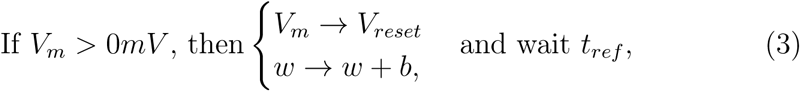

where *V_th_*is the adaptive spike initiation threshold, *t_ref_* the refractory period, *C* the membrane capacitance, *I_bias_*the bias current, *V_m_*the membrane potential, *g_L_*the leak conductance, *E_L_*the resting membrane potential, !*T* the sharpness factor, *ω_w_* the adaptation time constant, *a* the sub-threshold adaptation conductance, *b* the spike-triggered adaptation, and *V_reset_*the reset potential (36).

In the NEural Simulation Tool (NEST) (14), synaptic conductances are calculated and added directly to the calculation of the membrane potential. The excitatory synaptic conductance is represented by *g_e_*(*t*), the excitatory reversal potential by *E_e_*, the inhibitory conductance by *g_i_*(*t*), and the inhibitory reversal potential by *E_i_*. Gaussian white noise is added to the bias current of individual neurons at each time step. The mean is 0pA and standard deviation is equal to the current bias provided to that neuron. Noise can be negative (inhibitory) or positive (excitatory). The exact values for constants can be found in Table in the Supplementary Material. The parameters are tuned to produce the desired firing behavior as well as a biologically plausible instantaneous spiking frequency. During in vivo walking, the majority of spinal neurons, including fast motoneurons, are firing at 50-100Hz (46; 22).

As rise and decay times of excitatory postsynaptic currents (EPSCs) and inhibitory postsynaptic currents (IPSCs) are affected during disease progression (35), we implemented a synaptic model where both of these time constants can be manipulated (visualization in Fig. in the Sup-plementary Material). We used NESTML (27) to create a version of the ’aeif_cond_alpha’ neuron model, substituting a beta function to control synaptic conductance. The excitatory and inhibitory dynamics of the beta function are shown in Equations 4.

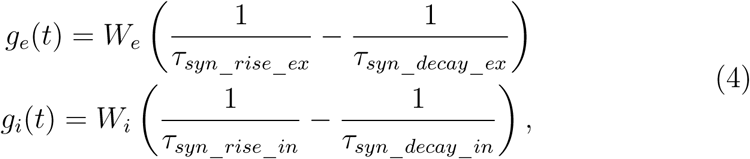

where *W_e_* represents the excitatory weight, *T_syn_*___*_rise_*___*_ex_* the excitatory rise time constant, *T_syn_*___*_decay_*___*_ex_* the excitatory decay time constant, *W_i_* the inhibitory weight, *T_syn_*___*_rise_*___*_in_* the inhibitory rise time constant, and *T_syn_*___*_decay_*___*_in_* the inhibitory decay time constant. The function is normalized so that if the weight is 1.0, the largest conductance will be 1nS. The synaptic weights are initialized as a distribution using a set mean and standard deviation depending on which populations are being connected. The strength of these connections was manually tuned based on network output and relevant synaptic weights were manipulated during disease progression as described in the Results section. Exact values for the synaptic weights can be found in the source code, available on GitHub (44).

### 2.2. Healthy Network

The computational model implemented in this study represents the flexor-extensor central pattern generator (CPG) of one side of the lumbar spinal cord (43; 24; 39; 6; 26; 40; 5) to produce the antagonistic output controlling the flexor-extensor phases of a single limb. Our model implements two rhythm-generating (RG) populations in a half-center oscillator architecture (17; 16), where the RGs mutually inhibit each other through two inhibitory interneuron populations (43; 40; 5; 24). Each RG comprises both excitatory and inhibitory neurons at a ratio of 4:1. Balance is manipulated through excitatory vs. inhibitory synapse strength (45). Furthermore, excitatory and inhibitory populations contain both bursting and tonically firing neurons (19). The excitatory neurons in the RGs approximate the Shox2 non-V2a, Hb9, and Lhx9 neurons, which have been linked to rhythm generation (10; 8). The number of neurons per population is given in Table in the Supplemen-tary Material. The connectivity details, including weights and percentages of connectivity between populations can be found in the source code on GitHub (set_network_params.py) (44).

The bursting neurons contribute to the generation of oscillatory activity in the RG population when receiving a tonic drive and the network frequency depends on the drive level, mimicking descending motor commands from brainstem locomotor centers. For each population receiving descending drive (see Fig. 1B), we applied a normally distributed input current to each neuron, with the population mean and a standard deviation of 25% of that mean (Table A.1). A tonic drive is also used to excite the Ia-afferent neuron populations to induce baseline firing in the populations. The flexor RG produced rhythmic activity when isolated from the network, while the extensor RG fired tonically when isolated (see Fig. 1C), following the state-dependent flexor-driven model for rhythm generation (19; 43; 6; 26). To produce oscillations in the flexor RG population, 30% of neurons were set to bursting, while the extensor RG included 10% bursting neurons to ensure tonic population activity.

We assert that the inhibitory populations connecting the RGs are equivalent to the V1*_RG_* and V2b inhibitory populations which are responsible for intralimb coordination (10) and are known to affect flexor/extensor alternation (47; 2). Downstream of the RGs, we included the interneuron populations known to be affected in the SOD1^G93A^ ALS mouse model (41; 34; 32). This included the excitatory interneuron populations V2a and V0c as well as the V1 inhibitory interneuron populations Ia and RC. Finally, motoneurons were implemented and acted as the output of the network. The number of neurons and firing behavior per population type are shown in Table in the Supplementary Material.

### 2.3. Degenerating Network

The network was validated at three progressive time points where the transcriptomic data from the SOD1^G93A^ mouse model provided information about degenerated cells (32). The number of remaining healthy cells across time points for each of the affected populations is outlined in Fig. 1E. Furthermore, all projections from V1 populations - V1*_RG_*, Ia, and RC were reduced by 50% from postnatal day 45 (P45) onward (2) (Fig. 1F). This was implemented through the percentage of connectivity, which describes the probability of a presynaptic neuron connecting to any postsynaptic partner. For example, the V1 subpopulations outside the RG layer projected at a rate of 28% to their postsynaptic partners in the healthy network, from P45 onwards, this is reduced to 14%.

### 2.4. Testing

The presented results show the variation across 25 trials. The same set of 25 seeds was used for each condition so that results were comparable. Several metrics were selected to compare network output: average maximum neuron firing rate, frequency, phase difference, and burst duration. The neuron firing rate plots are created by convolving individual spikes with a Gaussian window of 20ms. The highest peak was compared to the highest instantaneous firing rate for the population and then the convolved activity was adjusted to ensure the correct firing rate was recorded. This adjustment was calculated for each trial per population to ensure an accurate representation of the firing rate. For example, if the maximum instantaneous firing rate was calculated to be 100Hz and the peak of the convolved activity was 50, all values for the population in that trial would be multiplied by 2. The average maximum neuron firing rate was then calculated by averaging the peak values of the convolved plot per population within each trial. Frequency was calculated using Equations 5 and 6.

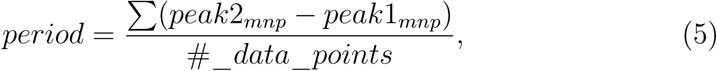

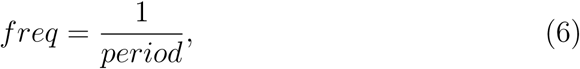

where *period* is the average period of either the flexor or extensor MNP over the simulation, *freq* is the average frequency of either the flexor or extensor MNP over the simulation, *avg*_*period* is the average period of both MNP signals over the simulation. The period and frequency of the signals were calculated using the peaks of the signals, *peak*2*_mnp_* and *peak*1*_mnp_*. When calculating using ’findpeaks’, the minimum peak height was set to 40% of the maximum amplitude of the signal, the minimum distance between peaks was set to 1000ms, and prominence was set to 0.1. Burst duration was calculated by recording when the population outputs crossed each other and subtracting the time between. The durations of the flexor and extensor MNP bursts were recorded separately. All burst durations within a single trial for the flexor and extensor were then averaged separately. Phase was determined by calculating the midpoint between the crossings for the flexor and extensor MNPs, as shown in Equation 7.

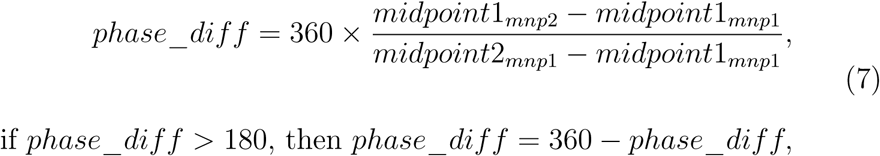

where *phase*_*diff* is the phase difference between flexor and extensor MNP outputs, *midpoint*1*_mnp_*_2_ is the midpoint of the current extensor burst, *midpoint*1*_mnp_*_1_ is the midpoint of the current flexor burst, and *midpoint*2*_mnp_*_1_ is the midpoint of the next flexor burst.

We used two statistical tests to compare the results depending on whether two or more conditions were compared. In all experiments, we tested 25 trials per condition and we excluded outlier trials that were outside 3 standard deviations. We applied the non-parametric Kruskal-Wallis H-test with Dunn’s post hoc test when comparing experiments. All disease time points were compared to the healthy control. When we compared flexor versus extensor, we applied a Wilcoxon signed-rank test. Our significance levels were defined as: p *<*= 0.001 = ***, 0.001 > p *<*= 0.01 = **, 0.01 > p *<*= 0.05: *, p > 0.05 = ns.

### Ethical permits and mouse strains

All experiments performed on mouse models comply with the relevant regulations of the EU Directive 2010/63/EU as well as the UK Home Office Guidance on the Operation of the Animals Act 1986. The gene therapy procedures for the delivery of systemic administration of AAVs were performed at University of Copenhagen and approved by the Danish Animal Experiments Inspectorate (Ethical permit: 2022-15-0201-01164). Tissue analysis was performed at University of St Andrews upon ethical approval of experimental methods obtained from the Animal Welfare Ethics Committee of the University of St. Andrews under the permit PS18278. SOD1^G93A^ mice (B6.Cg-Tg(SOD1-G93A)1Gur/J stock no: #004435) were obtained from the Jackson Laboratory and kept on a congenic background. The En1^cre^ mice, kept on the C57BL6/J background, were previously obtained from Dr. Jay Bikoff (St. Jude Children’s Research Hospital, Memphis, TN, USA). Mice were genotyped using DNA extracted from ear clipping and the following primers were used: for the SOD1G93A gene, 5’-CAT CAG CCC TAA TCC ATC TGA-3’ and 5’-CGCGAC TAA CAA TCA AAG TGA-3’, while for the En1cre gene, 5’-GAG ATT TGC TCC ACC AGA GC-3’ and 5’-AGG CAA ATT TTG GTG TAC GG-3’. Copy number of the mutated SOD1 gene was quantified by qPCR utilizing the primers 5’-GGG AAG CTG TTG TCC CAA G-3’ and 5’-CAA GGG GAG GTA AAA GAG AGC-3’ for the SOD1G93A gene. The qPCR reaction was performed as suggested by the supplier’s protocol. All mouse strains were bred with congenic C57BL6/J mice, stock no: #000664 (Jackson Laboratory). Mice were housed according to standard conditions with ad libitum feeding, constant access to water, and a 12:12h light/dark cycle. All mice, including multiple crossings, were genotyped, tested for copy number, and phenotype was assessed, including weekly weight. For survival experiments, the humane endpoint was defined as a weight loss of 15% and/or functional paralysis in both hind limbs together with inability to perform a righting test<20s. Samples from five female and four male mice were used in this study, all of them carrying the SOD1^G93A^ mutation. All mice were euthanized between 145-170 postnatal days, once the humane endpoint was reached.

#### 2.4.1. Adeno-associated viral vector administration

The AAV-PHP.eB-hSYN-DIO-hESYT1-W3SL and the AAV-PHP.eB-hSYN-eGFP-W3SL vectors were generated as described in (34). Viral expression was under control of the human Synapsin (SYN) promoter for long-term over-expression in neurons. The PHP.eB capsid was used to increase transduction efficiency in the central nervous system and blood-brain-barrier crossing ability since the virus was intravenously administered (Link to Method). The viruses were produced by the Verhaagen lab at Netherlands Institute for Neuroscience. The viral production protocol uses cells in suspension, rather than monolayer plated cells, which enables production of a titer in the 10^11^ range. Systemic delivery of *ESYT1* was cre-dependent and En1^cre^ mice were used to ensure V1-restricted overexpression. The En1^cre^ mice were crossed with the SOD1^G93A^ mice to allow for V1-restricted gene therapy expression in ALS mice. The AAV-PHP.eB-hSYN-eGFP-W3SL vector expressing enhanced green fluorescent protein (eGFP) was used as a control and delivered to age-matched SOD1^G93A^ mice. Both AAVs were injected between P28 and P35 at a volume of 100*µ*L per mouse and a titer of 3.03×10^11^ viral genomes (vg) per 100*µ*L was used. Intravenous (IV) injections of AAVs were performed under general anesthesia with isoflurane (4% in 100% oxygen for induction, 1.5–2% in 100% oxygen for maintenance).

### Immunohistochemistry

To collect tissue for immunohistochemistry, mice were injected with Pentobarbital (250 mg/kg) and transcardially perfused with pre-chilled phosphate buffered saline (PBS, #10388739, Gibco) followed by pre-chilled 4% paraformaldehyde (PFA, #HL96753 HistoLab). The spinal cord was then dissected, postfixed for 60 min in cold 4% PFA and cryoprotected in PBS 30% sucrose for 48h at 4°C. Coronal sections of the lumbar spinal cord were sectioned at 30*µ*m-thickness and collected on Superfrost Plus slides (#12312148, Epredia). Slides were stored at –20°C until further processing. Slides were then washed for 10 min in PBS at room temperature. The samples were then incubated in primary antibodies overnight, using either a sheep anti-Chx10 (1:200, Millipore #AB9014), which is a marker for V2a INs, or guinea pig anti-VGlut2 (1:500, SYSY Antibodies #135 404), a marker for the glutamate transporter. Samples were then washed twice with PBS for five minutes each at room temperature and then incubated with secondary antibodies for an additional hour. The VGlut2 antibody was visualized with AlexaFluor 647 goat anti-guinea pig antibody (1:500, Thermo Fischer Scientific). The Chx10 antibody was visualized with AlexaFluor 488 donkey anti-sheep antibodies (1:500, Thermo Fisher Scientific). Two more PBS washes then took place for five minutes each. A Fluoro Nissl Neuro-Trace 530 counterstain (1:500, Thermo Fischer Scientific) was then applied for an hour to stain all neurons, and two final PBS washes were done for five minutes each. A negative control slide was also prepared using the same protocol without the primary Chx10 antibody incubation and NeuroTrace staining. All samples were then cover slipped using Fluoromount (Sigma Aldrich) for image acquisition.

### Image Acquisition and Quantification

Microphotographs were obtained using an Axio Observer 7 Zeiss epifluorescent microscope. Six spinal cord sections were analyzed per mouse and photographs of ventral horns of each hemicord were taken using a Plan-Apochromat 20x objective – NA=0.8, zoom=0.5. Images were then exported as .czi files and quantifications were performed utilizing the Fiji software. For pixel area analysis, all images were converted to grayscale and the threshold was set within ± 1000 intensity units per pixel for each image analyzed per mouse. The measurements were limited to the threshold. Images with a higher background/noise ratio were excluded. Statistical analysis was conducted in RStudio version 2023.06.2+561 and the results were analyzed using t-tests to compare V2a cell counts and VGlut2 synaptic areas between the *ESYT1* treated and untreated groups.

## Consent for publication

All authors approved the submitted version of the manuscript and consented to publication.

## Data availability

All data that support the findings of this study are included within the article and supplementary material. Data sets can be reproduced using the publicly available code.

## Materials availability

Not applicable

## Code availability

Code used to perform this study is publicly available on GitHub (https://github.com/Allodi-Lab/cpg_modeling_als/) (44).

## Author Contributions

BS and IA developed the main idea of the paper. BS researched the biological neural network concepts and implemented the computational model with support from JA. RMR and SM provided data for the paper and commented on the manuscript. KG performed experiments on the mouse model and quantified data. The manuscript was written by BS with support from JA and IA. JA and BS created the figure panels for the paper. JA and IA provided theoretical and experimental neuroscientific knowledge and contributed equally as senior authors.

## Acknowledgments

The authors would like to thank Associate Professor Alexander Matthias Walter at University of Copenhagen for his input regarding synaptic dynamics; Professor Joost Verhaagen and Dr Kimberly Pietersz at the Netherlands Institute for Neuroscience for the generation of the AAVs used in the study. Additionally, we would like to thank Professor Ole Kiehn for his valuable input on an earlier version of the manuscript. Finally, we would like to thank Dr. Filipe Nascimento for his insights into synaptic changes during ALS progression.

