## Supplementary Material for "Spinal Circuit Mechanisms Constrain Therapeutic Windows for ALS Intervention"

---

[†] The second affiliation is the authors' current affiliation.

#### Appendix A. Network and Neuronal Characteristics

Table A.1: Characteristics per neuron population.

| **Population** | **# of neurons** | **Firing behavior** | **Drive (pA)** |
|---|---|---|---|
| RG Flx exc | 240 | Bursting | 450 ± 112 |
| RG Flx exc | 560 | Tonic | 720 ± 180 |
| RG Flx inh | 60 | Bursting | 450 ± 112 |
| RG Flx inh | 140 | Tonic | 720 ± 180 |
| RG Ext exc | 80 | Bursting | 840 ± 210 |
| RG Ext exc | 720 | Tonic | 1344 ± 336 |
| RG Ext inh | 20 | Bursting | 840 ± 210 |
| RG Ext inh | 180 | Tonic | 1344 ± 336 |
| $V1_{RG}$ | 300 | Tonic | 144 ± 36 |
| V2b | 300 | Tonic | 182 ± 45 |
| V2a | 158 | Tonic | 480 ± 120 |
| V0c | 15 | Tonic | 480 ± 120 |
| Ia | 60 | Tonic | 240 ± 60 |
| RC | 30 | Bursting | 0 |
| Motoneuron | 150 | Tonic | 0 |

Table A.2: Parameter values to produce tonic firing and bursting behavior. RG = Rhythm Generator, RC = Renshaw Cell

| **Parameter** | **Tonic firing** | **Bursting (RG)** | **Bursting (RC)** |
|---|---|---|---|
| $V_{th}$ (mV) | -50 ± 1 | -51 ± 1 | -51 ± 1 |
| $t_{ref}$ (ms) | 9 ± 0.2 | 3 ± 0.2 | 3 ± 0.2 |
| $C$ (pF) | 200 ± 40 | 600 ± 80 | 600 ± 80 |
| $g_L$ (nS) | 10 | 26 | 26 |
| $E_L$ (mV) | -70 | -60 | -60 |
| $\Delta T$ (mV) | 2 | 3 | 2 |
| $\tau_w$ (ms) | 30 | 260 | 130 |
| $a$ (nS) | 3 | -11 | -11 |
| $b$ (pA) | 0 | 60 | 30 |
| $V_{reset}$ (mV) | -58 | -48 | -48 |

### Appendix B. Statistics

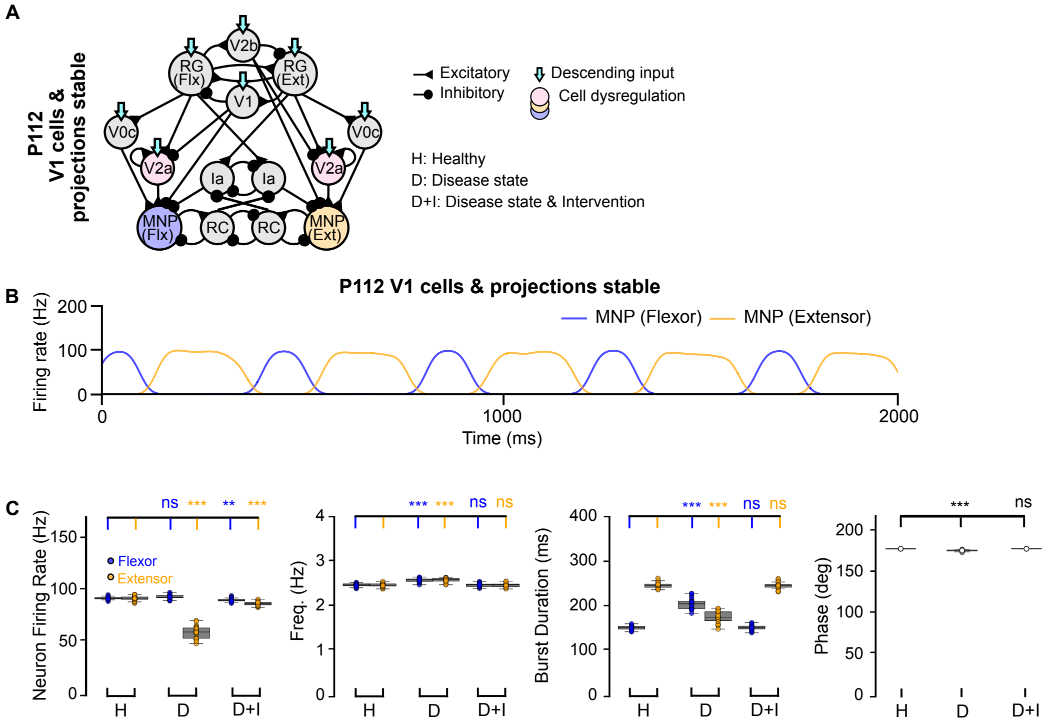


D

**Mean and standard deviation per metric under test conditions**

| Timepoint | % Outlier | Max IFR Flx | Max IFR Ext | Freq Flx | Freq Ext | BD Flx | BD Ext | MNP Phase |
|---|---|---|---|---|---|---|---|---|
| **V1 cells and projections stable (comparing to healthy and no intervention)** | | | | | | | | |
| P112 | 0 | 91.16±1.51 | 87.73±2.0 | 2.49±0.04 | 2.49±0.04 | 153.71±5.31 | 247.37±7.14 | 179.22±0.13 |
| p-value vs healthy | | 1.93e-03 | 3.41e-04 | 1.00e00 | 1.00e00 | 1.00e00 | 1.00e00 | 9.42e-01 |
| p-value flx vs ext | | 6.56e-06 | | 3.97e-01 | | 5.96e-08 | | |

Figure B.8: **Computational model of the locomotor CPG with V1 cells and projections stable but V2a interneuron and motoneuron loss. A**) Block diagram visualizing network changes. **B**) Example population output of the Flexor and Extensor motoneurons. **C**) Variance plots comparing neuron firing rate, frequency, burst duration, and phase. **D**) Statistics overview at P112 when V1 cells and projections are stable showing mean and standard deviation. P-values either compare the time point to healthy or flexor versus extensor within the time point. Significance across conditions was assessed with a Kruskal-Wallis H-test with Dunn's test post hoc, P-values comparing flexor versus extensor are calculated with a Wilcoxon Signed-Rank test, N=25, p <= 0.001 = ***, 0.001 > p <= 0.01 = **, 0.01 > p <= 0.05: *, p > 0.05 = ns.

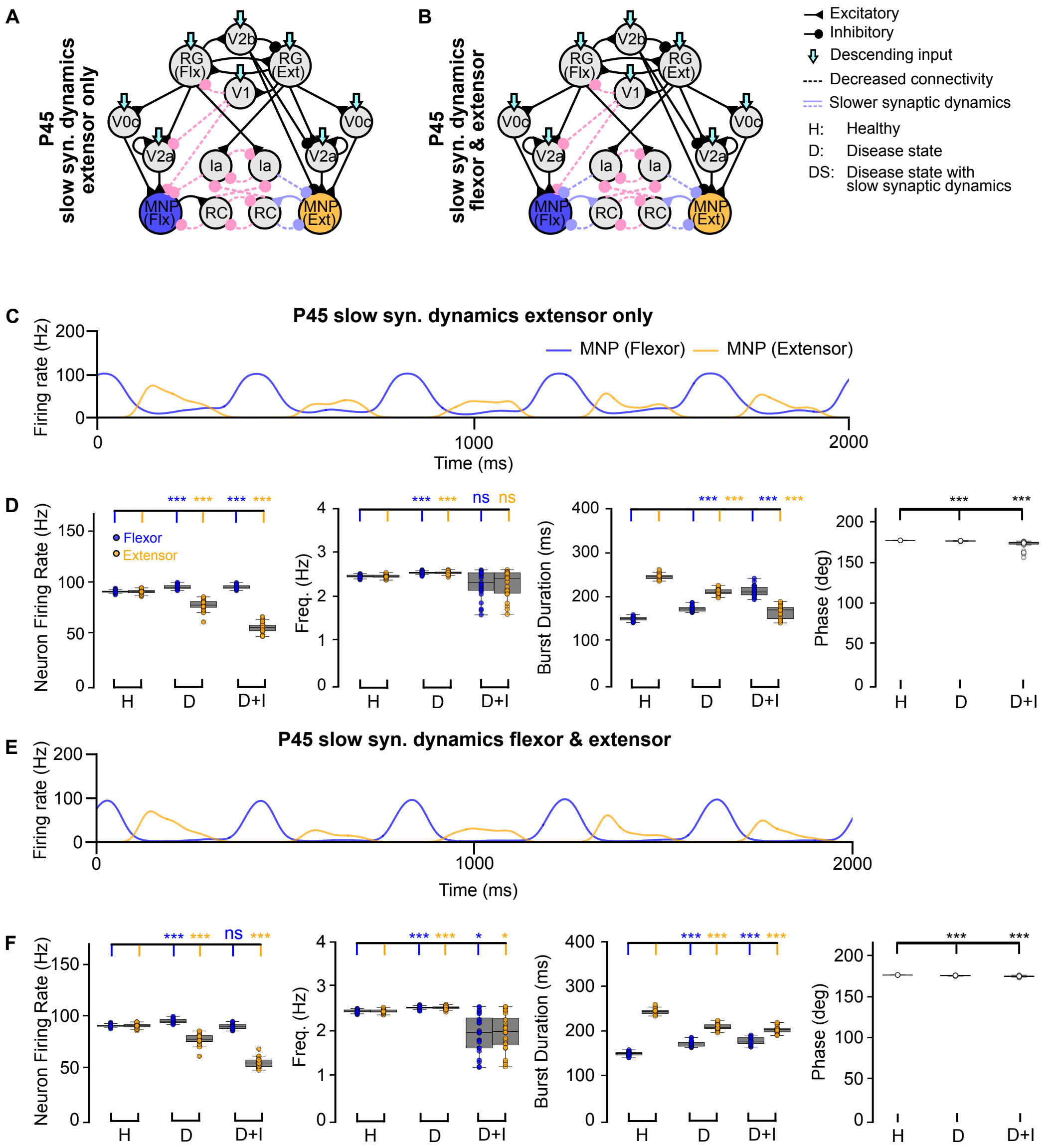


**G**

**Mean and standard deviation per metric under test conditions**

| Timepoint | % Outlier | Max IFR Flx | Max IFR Ext | Freq Flx | Freq Ext | BD Flx | BD Ext | MNP Phase |
|---|---|---|---|---|---|---|---|---|
| **Increased rise and decay times to RCs and MNs - extensor side only (comparing to healthy and no intervention)** | | | | | | | | |
| P45 | 0 | 97.48±1.95 | 58.2±4.88 | 2.31±0.3 | 2.31±0.03 | 215.5±11.63 | 169.38±14.9 | 173.73±5.5 |
| p-value vs healthy | | 1.70e-08 | 3.10e-15 | 1.00e00 | 1.00e00 | 2.37e-15 | 2.37e-15 | 3.82e-15 |
| p-value flx vs ext | | 5.96e-08 | | 1.00e00 | | 5.96e-08 | | |
| **Increased rise and decay times to RCs and MNs - flexor and extensor (comparing to healthy and no intervention)** | | | | | | | | |
| P45 | 4 | 92.06±2.65 | 57.31±4.47 | 1.99±0.42 | 2.0±0.41 | 181.57±7.77 | 207.9±7.47 | 177.64±0.66 |
| p-value vs healthy | | 1.00e00 | 4.21e-15 | 1.29e-02 | 2.38e-02 | 1.49e-11 | 1.23e-11 | 4.21e-13 |
| p-value flx vs ext | | 1.19e-07 | | 8.64e-01 | | 1.19e-07 | | |

Figure B.9: **Computational models of the locomotor CPG with slow synaptic dynamics. A,B**) Block diagram visualizing network changes. **C,E**) Example population output of the Flexor and Extensor motoneurons **D,F**) Variance plots comparing neuron firing rate, frequency, burst duration, and phase. **G**) Statistics overview at P45 during slow synaptic dynamics showing mean and standard deviation. P-values either compare the time point to healthy or flexor versus extensor within the time point. Significance annotation is calculated with a Kruskal-Wallis H-test with Dunn's test post hoc, P-values comparing flexor versus extensor are calculated with a Wilcoxon Signed-Rank test, N=25, p <= 0.001 = ***, 0.001 > p <= 0.01 = **, 0.01 > p <= 0.05: *, p > 0.05 = ns.

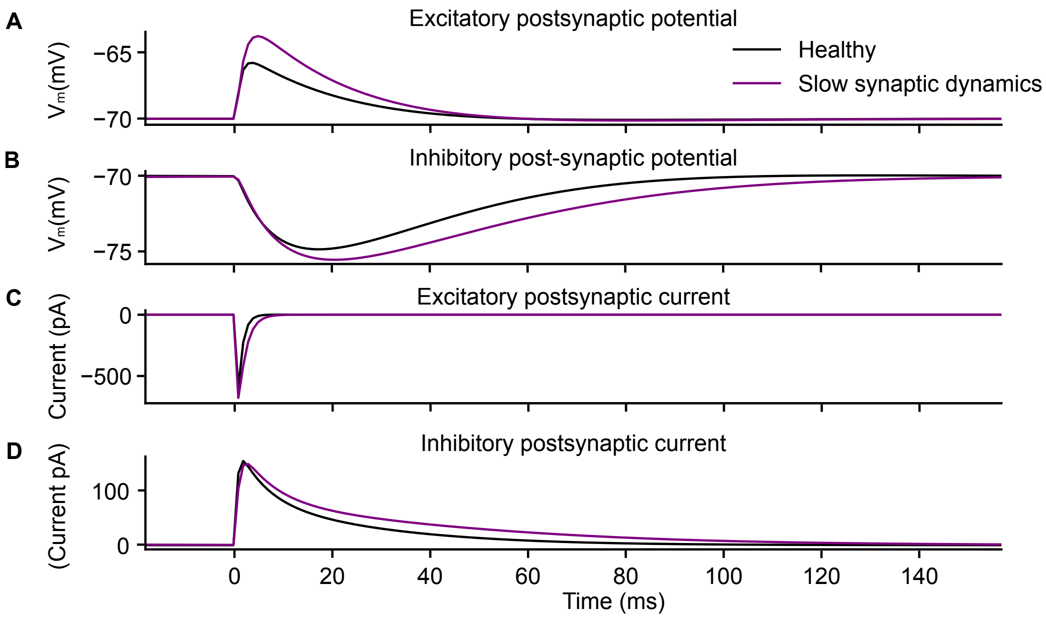


Figure B.10: **Comparison of healthy to slow synaptic dynamics.** Healthy (black) versus slow (purple) postsynaptic kinetics for excitatory and inhibitory synapses. **A**) Excitatory postsynaptic potential. **B**) Inhibitory postsynaptic potential. **C**) Excitatory postsynaptic current. **D**) Inhibitory postsynaptic current.

**A**

**Mean and standard deviation per metric at progressive timepoints**

| Timepoint | % Outlier | Max IFR Flx | Max IFR Ext | Freq Flx | Freq Ext | BD Flx | BD Ext | MNP Phase |
|---|---|---|---|---|---|---|---|---|
| Healthy | 0 | 92.91±1.25 | 92.93±2.09 | 2.49±0.03 | 2.49±0.04 | 153.29±4.49 | 248.35±6.14 | 179.18±0.12 |
| p-value flx vs ext | | 9.16e-01 | | 6.15e-01 | | 5.96e-08 | | |
| **No intervention** | | | | | | | | |
| P45 | 4 | 97.52±2.03 | 79.75±5.27 | 2.57±0.03 | 2.57±0.04 | 175.67±5.98 | 213.86±6.72 | 178.52±0.35 |
| p-value flx vs ext | | 1.19e-07 | | 8.86e-01 | | 1.19e-07 | | |
| P63 | 8 | 95.85±2.16 | 82.93±4.00 | 2.59±0.03 | 2.59±0.03 | 263.72±35.35 | 122.15±24.99 | 165.49±10.27 |
| p-value flx vs ext | | 2.38e-07 | | 8.91e-01 | | 2.38e-07 | | |
| P112 | 8 | 94.24±1.93 | 60.43±5.87 | 2.60±0.04 | 2.60±0.04 | 206.86±11.86 | 177.23±13.14 | 177.14±0.93 |
| p-value flx vs ext | | 2.38e-07 | | 6.57e-02 | | 1.03e-05 | | |

**B**

**Mean and standard deviation per metric under test conditions**

| Timepoint | % Outlier | Max IFR Flx | Max IFR Ext | Freq Flx | Freq Ext | BD Flx | BD Ext | MNP Phase |
|---|---|---|---|---|---|---|---|---|
| **No intervention (comparing to healthy and no intervention)** | | | | | | | | |
| MNs stable | 8 | 95.66±2.72 | 62.76±6.29 | 2.61±0.04 | 2.61±0.04 | 203.85±10.7 | 180.08±12.98 | 177.32±0.71 |
| p-value vs healthy | | 2.00e-05 | 1.89e-07 | 1.15e-08 | 3.80e-08 | 2.69e-08 | 3.27e-08 | 1.46e-08 |
| V2a stable | 4 | 99.54±1.83 | 85.81±3.5 | 2.61±0.04 | 2.61±0.04 | 344.04±97.69 | 92.84±19.21 | 146.0±17.23 |
| p-value vs healthy | | 1.94e-11 | 2.99e-04 | 2.00e-09 | 1.51e-08 | 5.96e-15 | 5.96e-15 | 6.64e-15 |
| V2a & MNs stable | 4 | 99.92±1.99 | 86.24±3.69 | 2.60±0.03 | 2.61±0.03 | 325.95±82.47 | 102.4±21.1 | 152.62±17.1 |
| p-value vs healthy | | 1.52e-11 | 6.12e-04 | 8.53e-09 | 2.40e-08 | 6.67e-15 | 5.96e-15 | 6.64e-15 |

**C**

**Mean and standard deviation per metric under test conditions**

| Timepoint | % Outlier | Max IFR Flx | Max IFR Ext | Freq Flx | Freq Ext | BD Flx | BD Ext | MNP Phase |
|---|---|---|---|---|---|---|---|---|
| **Exogenous synaptic stabilization (comparing to healthy and no intervention)** | | | | | | | | |
| P45 | 4 | 89.92±1.80 | 88.30±3.84 | 2.49±0.03 | 2.49±0.03 | 151.06±4.93 | 250.08±5.16 | 179.19±0.16 |
| p-value vs healthy | | 2.29e-03 | 2.34e-03 | 1.00e00 | 1.00e00 | 5.54e-01 | 1.00e00 | 1.00e00 |
| P63 | 0 | 91.58±2.04 | 86.74±5.09 | 2.47±0.04 | 2.47±0.04 | 159.08±6.54 | 246.31±6.14 | 179.10±0.22 |
| p-value vs healthy | | 2.19e-01 | 1.44e-04 | 3.20e-01 | 2.99e-01 | 1.32e-01 | 1.00e00 | 1.00e00 |
| P112 | 12 | 89.94±2.33 | 84.58±2.65 | 2.48±0.04 | 2.48±0.04 | 169.63±9.10 | 233.71±11.14 | 179.27±0.15 |
| p-value vs healthy | | 8.06e-04 | 2.97e-04 | 1.00e00 | 1.00e00 | 7.05e-04 | 7.67e-03 | 4.22e-01 |

**D**

**Mean and standard deviation per metric under test conditions**

| Timepoint | % Outlier | Max IFR Flx | Max IFR Ext | Freq Flx | Freq Ext | BD Flx | BD Ext | MNP Phase |
|---|---|---|---|---|---|---|---|---|
| **Exogenous synaptic stabilization (comparing to healthy and no intervention)** | | | | | | | | |
| MNs stable | 0 | 90.86±2.37 | 85.15±3.9 | 2.48±0.05 | 2.48±0.05 | 168.55±11.51 | 234.95±11.82 | 179.17±0.23 |
| p-value vs healthy | | 7.43e-03 | 1.85e-04 | 1.00e00 | 1.00e00 | 1.93e-03 | 1.74e-02 | 1.00e00 |
| V2a stable | 4 | 96.94±1.62 | 93.3±3.68 | 2.49±0.04 | 2.48±0.04 | 172.11±9.02 | 230.33±9.68 | 179.12±0.15 |
| p-value vs healthy | | 1.58e-09 | 1.00e00 | 1.00e00 | 1.00e00 | 3.95e-04 | 1.04e-03 | 1.00e00 |
| V2a & MNs stable | 0 | 97.05±2.2 | 95.12±3.11 | 2.48±0.04 | 2.48±0.04 | 168.04±11.23 | 235.2±11.78 | 179.16±0.18 |
| p-value vs healthy | | 1.78e-09 | 2.19e-01 | 1.00e00 | 1.00e00 | 4.43e-03 | 1.88e-02 | 1.00e00 |

| Timepoint | Max IFR |
|---|---|
| MNs stable | |
| p-value flx vs ext | 8.34e-07 |
| V2a stable | |
| p-value flx vs ext | 1.09e-03 |
| V2a & MNs stable | |
| p-value flx vs ext | 5.16e-02 |

**E**

**Mean and standard deviation per metric under test conditions**

| Timepoint | % Outlier | Max IFR Flx | Max IFR Ext | Freq Flx | Freq Ext | BD Flx | BD Ext | MNP Phase |
|---|---|---|---|---|---|---|---|---|
| **Increased rise and decay time to RCs and MNs (comparing to healthy and no intervention)** | | | | | | | | |
| P45 | 4 | 92.56±2.59 | 72.97±4.85 | 2.57±0.03 | 2.56±0.04 | 154.51±4.25 | 235.03±5.56 | 178.88±0.31 |
| p-value vs healthy | | 1.00e00 | 5.25e-13 | 2.44e-07 | 2.42e-06 | 1.00e00 | 6.02e-04 | 8.20e-03 |
| **Exogenous synaptic stabilization, Increased rise and decay times to RCs and MNs (comparing to healthy and no intervention)** | | | | | | | | |
| P45 | 4 | 76.38±5.75 | 88.18±4.01 | 2.49±0.03 | 2.49±0.04 | 139.51±4.53 | 261.59±5.13 | 179.18±0.14 |
| p-value vs healthy | | 1.14e-04 | 1.69e-03 | 1.00e00 | 1.00e00 | 4.14e-04 | 9.20e-04 | 1.00e00 |

Figure B.11: **Statistics per results panel.** **A**) Statistics overview at each time point without intervention showing mean and standard deviation. P-values compare flexor versus extensor within the time point. **B**) Statistics overview at P112 without and with synaptic stablization. P-values are in comparison to the healthy state. **C**) Statistics overview of each disease time point with synaptic stablization. P-values are in comparison to the healthy state. **D**) Statistics overview at P112 with synaptic stablization. P-values are in comparison to the healthy state as well as flexor versus extensor. **E**) Statistics overview at P45 with slow synaptic dynamics showing mean and standard deviation. P-values compare across conditions. P-values comparing across conditions are calculated with a Kruskal-Wallis H-test with Dunn's test post hoc, P-values comparing flexor versus extensor are calculated with a Wilcoxon Signed-Rank test, N=25.
